# Brain activation lateralization in monkeys (*Papio Anubis*) following asymmetric motor and auditory stimulations through functional Near Infrared Spectroscopy

**DOI:** 10.1101/2020.07.23.217760

**Authors:** C. Debracque, T. Gruber, R. Lacoste, D. Grandjean, A. Meguerditchian

## Abstract

Hemispheric asymmetries have long been seen as characterizing the human brain; yet, an increasing number of reports suggest the presence of such brain asymmetries in our closest primate relatives. However, most available data in non-human primates have so far been acquired as part of neurostructural approaches such as MRI, while comparative data in humans are often dynamically acquired as part of neurofunctional studies. In the present exploratory study in baboons (*Papio Anubis*), we tested whether brain lateralization could be recorded non-invasively using a functional Near-Infrared Spectroscopy (fNIRS) device in two contexts: motor and auditory passive stimulations. Under light propofol anaesthesia monitoring, three adult female baboons were exposed to a series of (1) left-*versus* right-arm passive movement stimulations; and (2) left-*versus* right-ear *versus* stereo auditory stimulations while recording fNIRS signals in the related brain areas (i.e., motor central sulcus and superior temporal cortices respectively). For the motor condition our results show that left-arm *versus* right-arm stimulations induced typical contralateral difference in hemispheric activation asymmetries in the three subjects for all three channels. For the auditory condition, we also revealed typical human-like patterns of hemispheric asymmetries in one subject for all three channels, namely (1) typical contralateral differences in hemispheric asymmetry between left-ear *versus* right-ear stimulations, and (2) a rightward asymmetry for stereo stimulations. Overall, our findings support the use of fNIRS to investigate brain processing in non-human primates from a functional perspective, opening the way for the development of non-invasive procedures in non-human primate brain research.

## Introduction

Lateralization is often presented as a key characteristic of the human brain, which separates it from other animal brains (1, 2); yet, an increasing number of studies, particularly in non-human primates (from here onward, primates), dispute this claim in a broad array of topics ranging from object manipulation, gestural communication to producing or listening to species-specific vocalizations (3-8). For instance, several primate studies present behavioral evidence of manual lateralization (4, 9), which have been associated with contralateral hemispheric correlates at the neurostructural level (5, 6). Other examples show orofacial asymmetries during vocal production, as evidenced by more pronounced grimaces on the left side of the mouth, which is suggestive of right hemisphere dominance in monkeys and great apes (7, 8), as has been documented in humans (10). In addition, comparative structural neuroimaging has shown that particular areas known to be leftwardly asymmetric in humans, such as the Planum Temporale in the temporal cortex, presented also leftward asymmetry in both monkeys and great apes (11-14), although the bias at the individual level seems more pronounced in humans (15, 16).

At the neural functional level using functional Magnetic Resonance Imaging (fMRI) or Positron Emission Tomography (PET) scan, most available studies in primates focused on lateralization of perception of synthesized sinusoidal or more complex vocal signals and reported inconsistent results. For instance, in rhesus macaques (*Macaca mulatta*), the processing of species-specific and/or heterospecific calls as well as non-vocal sounds, elicited various patterns of lateralized activations within the Superior Temporal Gyrus (STG) such as in the left lateral parabelt, either toward the right hemisphere or the left depending on the study (17-20). In chimpanzees (*Pan troglodytes*), a similar PET study reported a rightward activation within STG for processing conspecific calls (21). In general, such a variability of direction of hemispheric lateralization for processing calls appears similar to hemispheric lateralization variability described in humans for language processing depending of the type of auditory information and of language functions that are processed (22-24).

Compared to the leftward bias suggested for language, research investigating emotion perception in primates has strengthened the idea of a right bias in lateralization specific to emotion processing (3). For example, Parr and Hopkins (25) found that right ear temperature increased in captive chimpanzees when they were watching emotional videos, consistent with a greater right hemisphere involvement (25). The rightward hemisphere bias documented in chimpanzees is also found in other primate species such as olive baboons (*Papio anubis*) during natural interactions, as evidenced by studies investigating the perception of visual emotional stimuli (26-29). Yet, while the right hemisphere has understandably received much focused, the left hemisphere is also involved for emotion processing. For example, Schirmer and Kotz have suggested that the left hemisphere is particularly involved in the processing of short segmental information during emotional prosody decoding (24). Whether this functional differentiation, essential for speech perception in humans (30), is also present in non-humans is unclear. Baboons appear in this respect a particularly interesting animal model to study for lateralization, with several recent studies underlying the similarities in manual and brain asymmetries with humans (5, 14, 31). Furthermore, the baboon brain is on average twice as large as the macaque brain (32), which may facilitate the specific investigation of sensory regions. Finally, this species has all the primary cortical structures found in humans (33).

However, a major drawback in current studies lies in the complexity with which brain asymmetry can be investigated comparatively in primates. Here, we used functional Near-Infrared Spectroscopy (fNIRS) to test whether the blood oxygen level dependent (BOLD) response in baboon brains differed accordingly between the two hemispheres following left-*versus* right-asymmetric auditory and motor stimulations. fNIRS is a non-invasive optical imaging technique that has been developed to investigate brain processes in potentially at-risk populations such as human premature newborns, but which is now widely used with adult human participants. fNIRS is a relatively young imaging technique, with around two decades of use for functional research (34). Considering its portability and its lessened sensitivity to motion artefacts (35) compared to other non-invasive techniques, it might be an excellent methodology to study brain activations in primates under more ecologically relevant testing conditions, for example with a wireless and wearable device. As a first step, the present study tested fNIRS in baboons immobilized under light anesthesia monitoring. In relation with each of the stimulation types, we targeted relevant corresponding brain regions of interest – the motor cortex within the central sulcus and the auditory cortex regions in the temporal lobe respectively - by positioning the two sets of fNIRS channels in both hemispheres (one by hemisphere for a given region). We predicted that, if fNIRS was suitable to record brain signal in baboons, it would reflect contralateral hemispheric asymmetries in signals for each stimulation type within their corresponding brain region of interest, namely the motor cortex, associated with right- *versus* left-arm movements, and the temporal cortex, associated with the right- *versus* left- *versus* stereo ear auditory presentations. Our latter prediction was modulated by the knowledge that auditory regions are less lateralized, with about fifty percent of fibers projecting in the bilateral regions (36, 37), compared to cortical motor regions.

## Material & Methods

### Subjects

We tested three healthy female baboons (Talma, Rubis and Chet, mean age = 14.6 years, SD ±3.5 years). The subjects had normal hearing abilities and did not present a neurological impairment. All animal procedures were approved by the “C2EA -71 Ethical Committee of neurosciences” (INT Marseille) under the application number APAFIS#13553-201802151547729 v4, and were conducted at the Station de Primatologie CNRS (UPS 846, Rousset-Sur-Arc, France) within the agreement number C130877 for conducting experiments on vertebrate animals. All methods were performed in accordance with the relevant French law, CNRS guidelines and the European Union regulations (Directive 2010/63/EU). All monkeys were born in captivity from 1 (F1) or 2 generations (F2), and are housed in social groups at the Station de Primatologie in which they have free access to both outdoor and indoor areas. All enclosures are enriched by wooden and metallic climbing structures as well as substrate on the group to favour foraging behaviours. Water is available ad libitum and monkey pellets, seeds, fresh fruits and vegetables were given every day.

### Subject’s hand preference in communicative gesture and bi-manual task

The impacts of subject’s handedness on cerebral lateralization of language, motor and visual functions are well known in human neuroscience (38). For that purpose, we report here the hand preference of each baboon during visual communicative gesturing (CG - slapping one hand repetitively on the ground in the direction of a conspecific to threaten it) and during a bi-manual tube task (BM - holding a PVC tube with one hand while removing the food inside the tube with the fingers of the other hand). In both contexts, Talma was left-handed (CG: n=27, HI=-0.56, z-score=-2.89; BM: n=31, HI=-0.42, z-score=-2.33) whereas Rubis showed a preference toward the right hand (CG: n=16, HI=0.25, z-score = 1; BM: n=79, HI= 1, z-score=8.88). Conversely, Chet was left-handed in communicative gesture (n=25, HI = -0.44, z-score = -2.2) but right-handed in the bi-manual tube task (n=11, HI = 0.45, z=score = 1.51).

### Recordings

We selected one of the most wearable, wireless and light fNIRS devices available on the market (Portalite, Artinis Medical Systems B.V., Elst, The Netherlands) to measure the brain activations in baboons during the motor and auditory stimulations. The data were obtained at 50 Hz using six channels (three by hemisphere), three inter-distance probes (3 – 3.5 – 4 cm) and two wavelengths (760 and 850 nm). To localize our regions of interests (ROIs), the motor and auditory cortices, the fNIRS probes were placed using T1 MRI scanner images previously acquired by the LPC group on baboons (see Figure 1).

**Figure 1:**
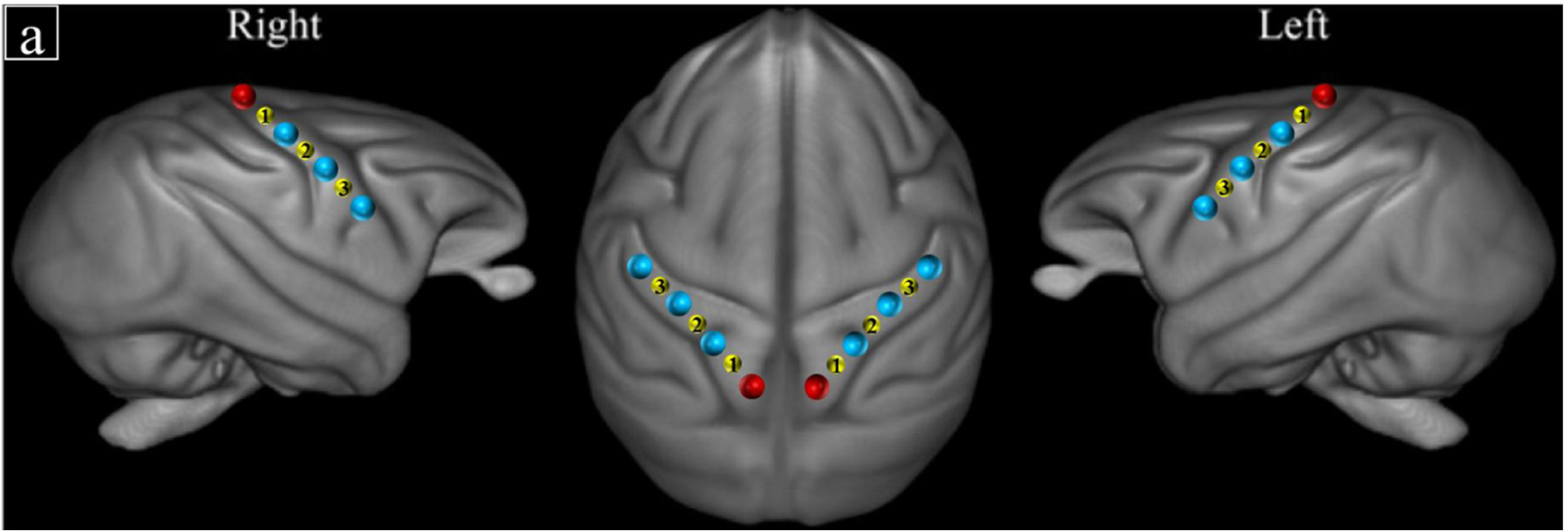
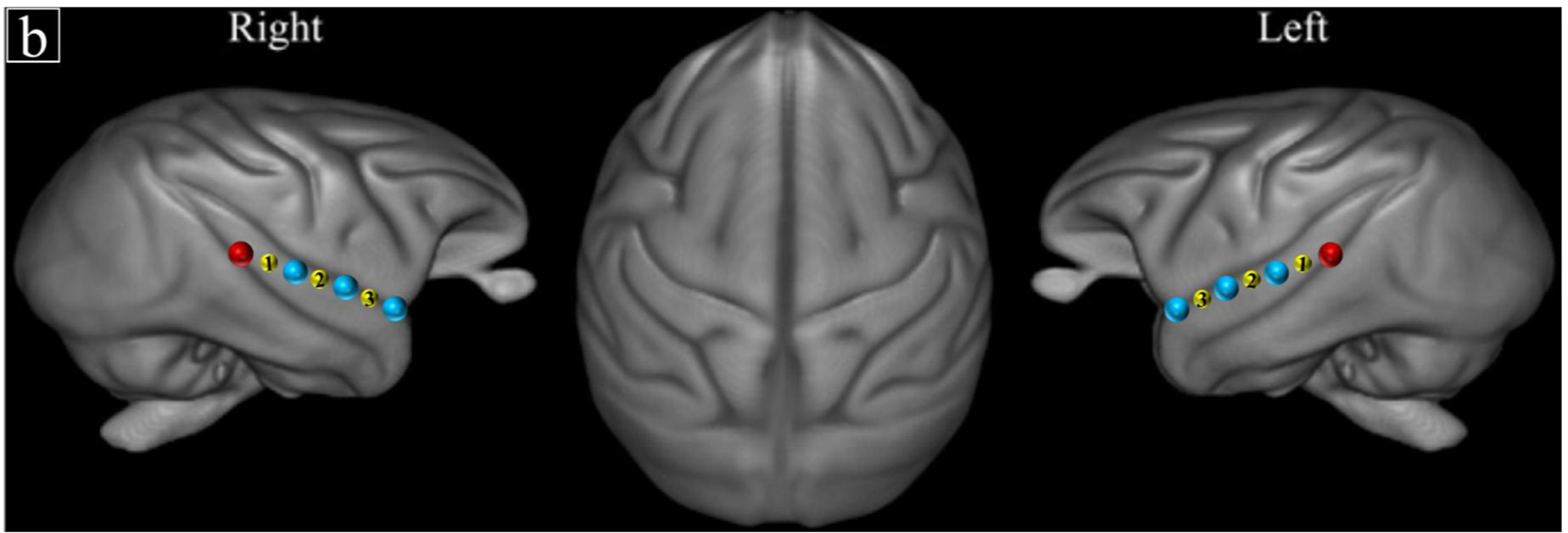
Schematic representation of fNIRS channel locations on ROIs according to T1 MRI template from 89 baboons (39) for (a) the motor and (b) the auditory stimulations. Red and blue dots indicate receivers and transmitters’ positions respectively. Yellow dots indicate the channel numbers.

Each fNIRS session was planned during a routine health inspection undergone by the baboons at the Station de Primatologie. As part of the health check, subjects were isolated from their social group and anesthetized with an intramuscular injection of ketamine (5 mg/kg - Ketamine 1000®) and medetomidine (50µg/kg - Domitor®). Then Sevoflurane (Sevotek®) at 3 to 5% and atipamezole (250 µg/kg - Antisedan®) were administered before recordings. The area of interest on the scalp was shaved. Each baboon was placed in *ventral decubitus* position on the table and the head of the individual was maintained using foam positioners, cushions and Velcro strips to remain straight and to reduce potential motion occurrences. Vital functions were monitored (SpO2, Respiratory rate, ECG, EtCO2, T°) and a drip of NaCl was put in place during the entire anaesthesia. Just before recording brain activations, sevoflurane inhalation was stopped and the focal subject was further sedated with a minimal amount of intravenous injection of Propofol (Propovet®) with a bolus of around 2mg/kg every 10 to 15 minutes or by infusion rate of 0.1 – 0.4 mg/kg/min. After the recovery period, baboons were put back in their social group at the Station de Primatologie and monitored by the veterinary staff.

### Motor stimulations

The motor stimulations consisted of 20 successive extensions of the same arm, alternatively right and left repeated three times according to the same set plan (L-R-R-L-L-R) for all baboons, resulting in a total of 120 arm movements. One experimenter on each side of the baboon extended slowly their respective arm while stimulating the interior side of the hand (gentle rhythmic tapping) with their fingers throughout the duration of the extension (about 5s) upon a brief vocal command triggered by another experimenter. Between each block, there was a 10s lag.

### Auditory stimulations

The auditory stimuli consisted of 20s-long series of agonistic vocalizations of baboons and of chimpanzees recorded in social settings (in captivity in an outside enclosure for baboons; and in the wild for chimpanzees). Equivalent white noise stimuli matched for the energy dynamics (i.e. the sound envelopes) were produced and used for comparison to control for the sound energy dynamic differences. In the present study and analysis, we only examine the effect of the lateralization of auditory stimulations (i.e., left ear *versus* right ear *versus stereo)* as a whole on hemispheric asymmetry and thus do not distinguish between auditory signal types or species (e.g. white noise and vocalizations). The auditory stimuli were broadcast pseudo-randomly, alternating voiced and white noise stimuli and separated by 15s silences, either binaurally (stereo), only on the left side, or only on the ride side. Due to signal artefacts and anaesthesia shortfalls, the number of stimuli between the three baboons differs slightly. For Talma, the total sequence consisted of 37 stimuli; for Rubis, the total sequence consisted of 47 stimuli; and for Chet, the total sequence consisted of 25 stimuli.

### fNIRS signal

We performed the first level analysis with MatLab 2018b (Mathwortks, Natick, MA) using the SPM_fNIRS toolbox (40, https://www.nitrc.org/projects/spm_fnirs/) and homemade scripts. Hemoglobin conversion and temporal preprocessing of O2Hb and HHb were made using the following procedure:

1. Hemoglobin concentration changes were calculated with the modified Beer-Lambert law (41);
2. Motion artifacts were removed manually in each individual and each channel for the auditory stimulations. Thus, 10 seconds in total (1.3%) were removed from the O_2_Hb and HHb signals of Rubis and 35 seconds (4.8%) for Talma and Chet fNIRS data;
3. A low-pass filter based on the hemodynamic response function (HRF) (42) was applied to reduce physiological confounds.
4. A baseline correction was used for both the motor and auditory stimulations by subtracting respectively (i) the average of 10 seconds intervals preceding each block; (ii) the average of the 15 seconds of silence preceding each sound.

According to the temporal properties of the BOLD responses for each baboon, the O2Hb concentration was averaged for Talma in a window of 4 to 12 s post stimulus onset for each trial; and for Rubis and Chet in a window of 2 to 8 s post stimulus onset in order to select the range of maximum concentration changes (µM). The difference of concentration range is explained by the presence of some tachycardiac episodes for both Rubis and Chet during the experiment, involving an HRF almost twice as fast as the one found for Talma.

### AQ score calculation

Asymmetry Quotients (AQ) were derived for each subject and each experimental condition (i.e: stimulation of the right arm and of the left arm for the motor experiment; right, left and stereo audio stimulation for the auditory blocks) by first calculating the difference between the right hemisphere (RH) and the left hemisphere (LH) values, to which we subsequently subtracted the same difference during the preceding baseline block for the same subject to normalize across trials. In particular, for motor stimuli, the baseline represented the 10s block without motor activity immediately before a passive stimulation block of the right or left arm. For auditory stimuli, the baseline was calculated on the 15s silence block that immediately preceded the auditory stimuli. In this analysis, all auditory stimuli (baboon and chimpanzee calls, and corresponding white noises) were analysed together. All calculated AQs were then normalized using the scale function of R studio (R studio (2015) Inc:, Boston, MA, url: http://www.rstudio.com/). For this analysis, we excluded one block ‘chimpanzee white noise audio stereo’ (2.7% of O2Hb signal) for Rubis, and two blocks ‘chimpanzee white noise audio stereo’ and ‘baboon white noise audio stereo’ (8.3%) for Talma as the recorded data revealed themselves artefactual beyond repair. Positive AQ values indicate a rightward asymmetry and negative values indicate a leftward asymmetry. Finally, using the aov function of R studio, we performed one-way ANOVAs with pairwise comparisons on individual baboons by comparing the AQ of all trials in the different stimulation conditions (right *versus* left motor stimulation; right *versus* left *versus* stereo auditory stimulation) enabling to generalize the data of each individual.

## Results

### Motor stimulations

One-way Anova analyses revealed significant differences between the left and right arm stimulations across the three channels and baboons. Hence, for Rubis and Chet, comparisons between left and right arms stimulations were all significant at p < .001 (Rubis: Ch1: *F*_1,118_ = 52.63; Ch2: *F*_1,118_ = 50.63; and Ch3: *F*_1,118_ = 42.35; for Chet: Ch1: *F*_1,118_ = 30.16; Ch2: *F*_1,118_ = 28.21; and Ch3: *F*_1,118_ = 24.77). Regarding Talma, significant differences were found at p <.05 in channel 1 (*F*_1,118_ = 3.821) and channel 3 (*F*_1,118_ = 6.521). The pairwise comparison in channel 2 (*F*_1,118_ = 14.71) was significant at p < .001.

Overall, the difference of AQ between left- *versus* right-arm stimulations were consistently contralateral across the three subjects for all three channels: activation asymmetries were more leftward for right-arm stimulations than for left arm stimulations and, were more rightward for left-arm stimulations than for right arm stimulations (Figure 2; see Table 1 in supplementary material for the mean AQ values).

**Figure 2:**
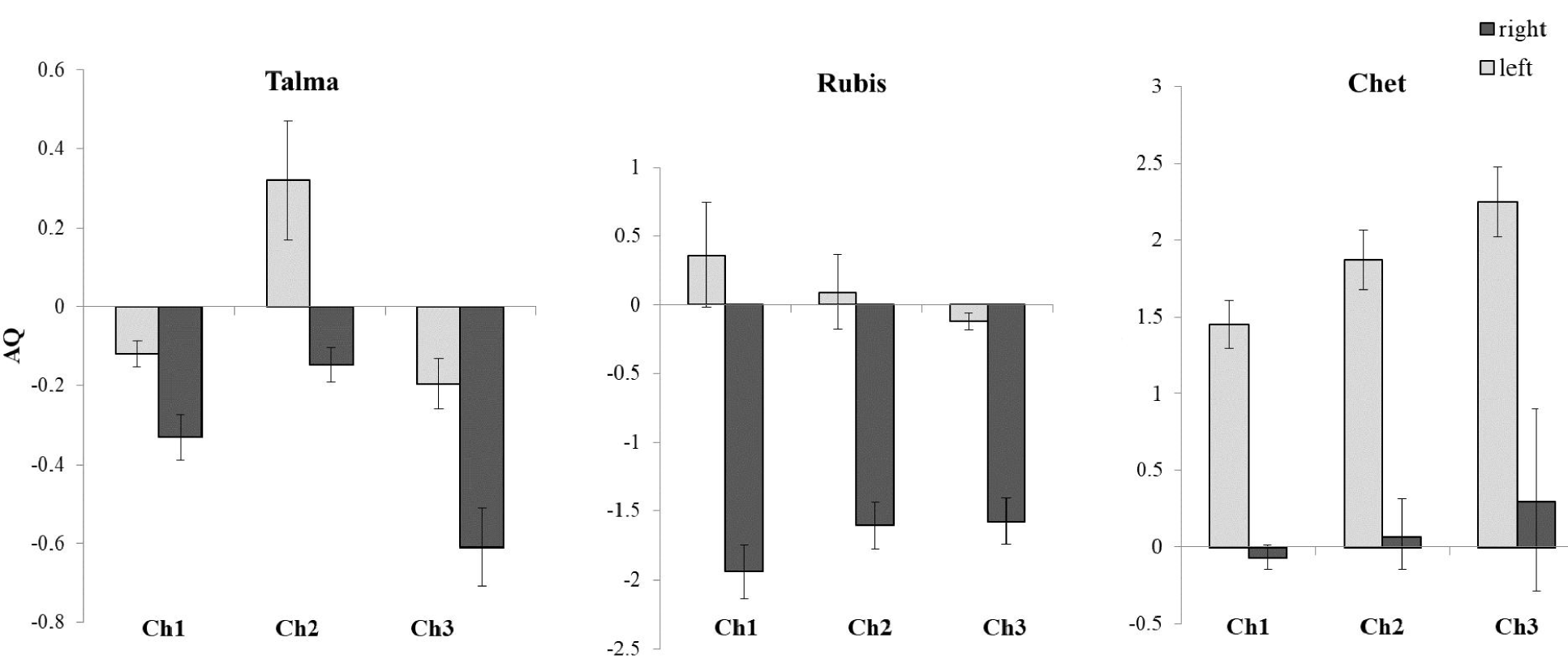
Normalized averaged AQ (and corresponding SE) in the motor cortex following motor stimulations in the three adult female baboons (see Figure 1 for localization of the channels).

### Auditory stimulations

We only found significant overall differences between, right, left and stereo ear stimulations (p <.05) for subject Chet (Figure 3) for all channels (Ch1: *F*_2,6_ = 7.073; Ch2: *F*_2,6_ = 6.473; and Ch3: *F*_2,6_ = 4.289). Pairwise comparison for right *versus* left ear stimulations were significant (p <.05) in Ch1 (*F*_1,6_ = 5.216) and Ch2 (*F*_1,6_ = 5.043). Furthermore, significant differences between right and stereo ear stimulations appeared across all channels (Ch1: *F*_1,6_ = 22.55; Ch2: *F*_1,6_ = 16.56, p <.001; Ch3: *F*_1,6_ =15.95, p <.05). Note that the comparison left *versus* stereo did not reach significance for any channels (Ch1: *F*_1,6_ = 1.827; Ch2: *F*_1,6_ = 1.825; Ch3: *F*_1,6_ =0.989, all p >.05).

**Figure 3:**
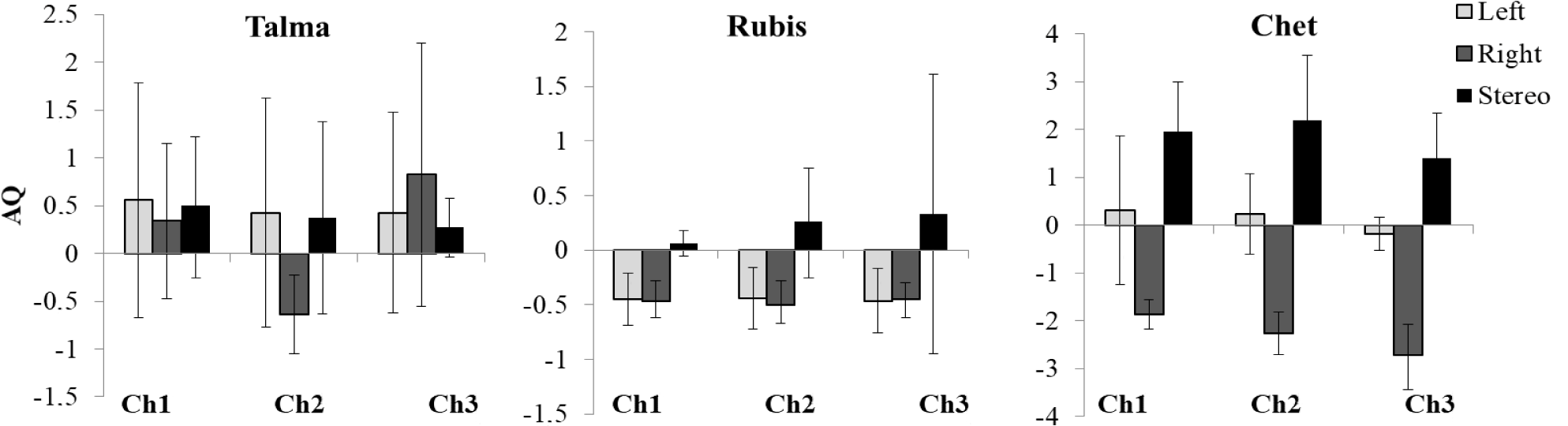
Normalized averaged AQ (and corresponding SE) above the temporal cortex following auditory stimulations in three adult female baboons (see Figure 1 for localization of the channels).

Hence, for Chet, there was a larger bias toward the left hemisphere with right ear stimulation compared to stereo (for all our channels) and left ear stimulation (for channels 1 and 2 only; Figure 3). No difference was recorded as significant for the two other baboons (see Table 2 in supplementary material for the mean AQ values).

## Discussion

The results of the present study clearly demonstrate that non-invasive fNIRS is a valid imaging technique to investigate functional lateralization paradigms in a nonhuman primate species.

Our most potent results were found with the motor stimulation where we observed a strong contralateral hemispheric asymmetry of the fNIRS signals in the motor cortex across baboons. Right arm movements elicited greater leftward asymmetry than left arm movements and *vice versa* in each of the three baboons for all three fNIRS channels. Results were clear-cut for Rubis and Chet, though interestingly opposed, with Rubis having a strong leftward asymmetry as a result of her right arm being stimulated, and Chet showing a strong rightward asymmetry for her left arm. Results for Talma were somewhat similar to Rubis’ since right arm movements elicited more leftward asymmetry than the left arm in channels 1 and 3. Results in channel 2 were most in line with our original prediction, namely a clear mirror pattern of contralateral asymmetries between the two arms: the right arm movements elicited leftward asymmetry and the left arm, a rightward asymmetry. Our results are consistent with previous studies in primates: for arm/hand movements, 90% of the corticospinal pathway project to the contralateral spinal cord (43-47). Hence, our study replicates these findings, with brain signals differences detected by non-invasive fNIRS. Despite the robust consistency of findings across subjects concerning the direction of the effect between the left and the right arms, the reasons for inter-individual variabilities as well as the lack of mirror pattern of results between the two arms (channel 2 of Talma excepted) remains unclear. In particular, potential involuntary differences in arms’ stimulation degree between the two experimenters involved in each of the subject’s arms manipulations, as well the handedness of each individual baboon may have had an impact on our results.

Our results were also consistent with predicted asymmetries regarding auditory stimulations for one subject. Contralateral differences of asymmetry were found for Chet in all three channels, with the stimulation of both ears and left ear eliciting overall more rightward asymmetries than right ear stimulations. Nevertheless, for Talma and Rubis, the direction and degree of asymmetry varied irrelevantly of whether the sound was presented to the right or left ear, namely toward the left temporal areas for Rubis and toward the right temporal areas for Talma. These mixed results related to auditory stimulation might be interpreted with respect to some characteristics of the hemispheric organization of the brain. It is well-known that at least one third of the auditory fibres from the olivary complex project to ipsilateral brain regions inducing less lateralization compared to motor brain regions. Furthermore, it has been shown that receptive fields in some regions sensitive to somatosensory input from the auditory cortex are 50% contralateral and 50% bilateral (48, 49); and that temporal regions such as the belt, parabelt and STS receive strong ipsilateral connections in rhesus macaques (50, 51), suggesting overall a less marked lateralization for auditory processing compared to motor regions. Interestingly, the subject’s handedness in communicative gesture could also explain these mixed results. In fact, our left-handed subject Talma, showed a clear right hemisphere bias for most stimuli (to the exception of the right ear stimulation in channel 2); whereas Rubis, right-handed in communicative gesture, showed a stronger bias toward the left hemisphere for the sounds broadcast in right and left ears. These preliminary findings may thus highlight the impact of hand preference in communicative contexts on contralateral brain organization in baboons during auditory processing but would need further investigations in a larger cohort of subjects.

Overall, given the lack of statistical power related to low sample size, we cannot draw any conclusion regarding the direction of hemispheric lateralizations at a population-level for sounds processing in baboons, or their relation to hand preference for communicative gesturing. Nevertheless, some of our findings remain consistent with the literature on human auditory pathways: for example, Kaiser and collaborators found that stimuli presented in stereo activated more the right hemisphere compared to lateralized sounds showing a left hemisphere bias (52). These results suggest that stereo sounds involve additional processing steps resulting in stronger and more rightward brain activations (53). This pattern of rightward asymmetry for stereo and left sounds processing in the baboon “Chet” is also somewhat consistent with previous rightward asymmetries reported in rhesus monkeys (17) and in chimpanzees (21) for processing conspecific calls. Hence, our data suggest that a phylogenetic functional approach to vocal perception appears possible with fNIRS.

In conclusion, our study shows that fNIRS is a valid methodology to access brain signals in primates non-invasively. In particular, we have replicated findings in the literature about brain contralateral hemispheric activation in two different modalities showing that fNIRS is able to capture such functional differences even in a context in which baboons were anesthetized. However, we have also uncovered large variation between individuals. This may be due to interindividual differences leading to the inability to precisely record in the same spot for all baboons. Indeed, while we based our placing of optodes on our subjects based on an averaged structural MRI pattern to which all tested individuals contributed, we cannot exclude small variation across cortices. In the future, fNIRS should thus be coupled with structural imaging techniques such as MRI that allow a precise positioning of the optodes for each individual. Yet, the need to couple fNIRS with existing techniques does not deny a more widespread use of fNIRS in the future. To the contrary, we believe that our study opens new avenues for brain investigation in nonhuman primates using fNIRS for two main reasons. First, fNIRS has been used in a multitude of contexts when other brain imaging techniques could not be used, for example in the field with greater ecological conditions (54). While our data have been recorded in anesthetized baboons, a logical next step is to train and habituate baboons to accept wearing a fNIRS device. Our experimental paradigms could then be extended in awake monkeys with more sophisticated design involving behavioural contingencies related to different kinds of stimulation. Second, our study stresses that fNIRS could in the future become a valuable method to explore brain activations in lateral regions in a non-invasive way in nonhuman animals without attempting the physical integrity of the subjects, which would ultimately make investigation of brain mechanisms in animal much more accessible and flexible.

## Acknowledgements

CD and TG were supported by the Swiss National Science Foundation (grants P1GEP1_181492 to CD and CR13I1_162720 / 1 to DG-TG). AM has received funding from the European Research Council under the European Union’s Horizon 2020 research and innovation program grant agreement No 716931 - GESTIMAGE - ERC-2016-STG. We thank the Société Académique de Genève for their financial support allowing purchasing the fNIRS equipment. We thank the vet Pascaline Boitelle for monitoring heath and anaesthesia of baboons during experiment and the animal care staff as well as Jeanne Caron-Guyon, Lola Rivoal, Théophane Piette and Jérémy Roche for assistance during the recordings.

## Notes

### Competing Interest Statement

The authors have declared no competing interest.

