## Supplementary material for "Brain activation lateralization in monkeys (*Papio Anubis*) following asymmetric motor and auditory stimulations through functional Near Infrared Spectroscopy"

Calculated mean AQ:

Asymmetry Quotients (AQ) were derived for each subject and each experimental condition (i.e: stimulation of the right arm and of the left arm for the motor experiment; right, left and stereo audio stimulation for the auditory blocks) by first calculating the difference between the right hemisphere (RH) and the left hemisphere (LH) values, to which we subsequently subtracted the same difference during the preceding baseline block for the same subject to normalize across trials. In particular, for motor stimuli, the baseline represented the 10s block without motor activity immediately before a passive stimulation block of the right or left arm. For auditory stimuli, the baseline was calculated on the 15s silence block that immediately preceded the auditory stimuli. In this analysis, all auditory stimuli (baboon and chimpanzee calls, and corresponding white noises) were analysed together. All calculated AQs were then normalized using the scale function of R studio (R studio (2015) Inc:, Boston, MA, url: http://www.rstudio.com/). For this analysis, we excluded one block 'chimpanzee white noise audio stereo' (2.7% of $O_2$Hb signal) for Rubis, and two blocks 'chimpanzee white noise audio stereo' and 'baboon white noise audio stereo' (8.3%) for Talma as the recorded data revealed themselves artefactual beyond repair. Positive AQ values indicate a rightward asymmetry and negative values indicate a leftward asymmetry.

Table 1: Calculated mean AQ values for subjects Talma, Rubis and Chet during the motor stimulation.

| | **Channel 1** | | **Channel 2** | | **Channel 3** | |
|---|---|---|---|---|---|---|
| | **Left** | **Right** | **Left** | **Right** | **Left** | **Right** |
| **Talma** | -0.119 | -0.330 | 0.320 | -0.146 | 0.195 | -0.609 |
| **Rubis** | 0.362 | -1.940 | 0.089 | 1.603 | -0.117 | -1.572 |
| **Chet** | 1.448 | -0.067 | 1.871 | 0.064 | 2.247 | 0.299 |

Table 2: Calculated mean AQ values for subjects Talma, Rubis and Chet during the auditory stimulation

| | **Channel 1** | | | **Channel 2** | | | **Channel 3** | | |
|---|---|---|---|---|---|---|---|---|---|
| | **Left** | **Right** | **Stereo** | **Left** | **Right** | **Stereo** | **Left** | **Right** | **Stereo** |
| **Talma** | 0.557 | 0.339 | 0.497 | 0.426 | -0.639 | 0.371 | 0.426 | 0.822 | 0.268 |
| **Rubis** | -0.448 | -0.468 | 0.060 | -0.438 | -0.501 | 0.258 | -0.462 | -0.449 | 0.332 |
| **Chet** | 0.312 | -1.865 | 1.961 | 0.226 | -2.268 | 2.189 | -0.185 | -2.718 | 1.395 |
